# Good moms: dependent-young and their mothers cope better than others with longer dry season in plains zebras

**DOI:** 10.1101/2021.10.18.464846

**Authors:** Lucie Thel, Christophe Bonenfant, Simon Chamaillé-Jammes

## Abstract

In large herbivores, the timing of births often coincides with the seasonal peak of food resources availability, likely to improve juvenile survival and reduce reproduction costs. Some species however, breed year-round, even in seasonal environments. Demographic processes, such as to what extent being born during the lean season reduces survival of juveniles and reproductive females, remain understudied in large mammals inhabiting tropical ecosystems. We investigated survival rates in plains zebras (*Equus quagga*) in Hwange National Park (Zimbabwe), a highly seasonal savanna ecosystem. We used capture-recapture models to analyse long-term demographic data (2008-2019). We investigated the effect of seasonality as a categorical (wet versus dry season) and continuous (duration of the dry season) variable on survival. We found little variability in early juvenile survival (φ = 0.458 ± 0.044 SE, < 6 m.o.), whereas late juvenile and yearling survival was higher and decreased with increasing length of the dry season (from 0.850 ± 0.095 SE to 0.480 ± 0.120 SE). Female survival was high (> 0.703 ± 0.057 SE and up to 0.995 ± 0.006 SE) but decreased with increasing exposure to the dry season in non-reproductive females. The probability of females becoming reproductive in the following year was not affected by the length of the dry season (0.423 and 0.420 for reproductive and non-reproductive females respectively). Our results highlight the importance of individual quality in reproductive performance, as reproductive females seem to buffer the effect of environmental variability on their own survival and that of their foal.

## Introduction

In long-lived species, the recruitment of juveniles in a population is often the most variable demographic rate and a major determinant of the variation in population growth rate (Gaillard et al. 1998, Gaillardet al. 2000, Kiffner and Lee 2019). Specifically, in large mammalian herbivores, recruitment depends mostly on juvenile survival (Gaillard et al. 2003), which is influenced primarily by mothers during the last months of gestation and by newborns during their first year of life (Lindström 1999, Beckerman et al. 2003, Kiffner and Lee 2019). For instance, exposure to harsh winters conditions reduces fawn survival in red deer (*Cervus elaphus*, Loison and Langvaten 1998). In reindeer (*Rangifer tarandus*), high rain-on-snow during the gestation period increases the negative effect of a low body mass of the mother on her probability to have a calf at heel during the next summer (Pigeon et al. 2022). For capital breeders, females usually enter a reproductive phase if they reach a threshold body mass (Jönsson 1997). The energy reserves built prior to reproduction critically contribute to the female’s ability to feed the fetus during pregnancy and the newborn during lactation (Stephens et al. 2009). Therefore, the level of reserves affects the weight of the newborn at birth (Cameron et al. 1993) and its survival to weaning (Jackson et al. 2021). In seasonal environments where food resource availability and quality vary strongly in a year, the timing of developmental stages and of the reproductive cycle is therefore critical for juvenile survival and the mother’s reproductive success.

In temperate and boreal ecosystems, the timing of birth contributes to silver-spoon effects on life history traits (*sensu* Grafen 1988). By determining the early environmental conditions experienced by newborns during the first months of life, timing of birth contributes directly (*e.g.* through thermal stress; Grovenburg et al. 2012) and indirectly (*e.g.* through potentially lower food availability reducing growth rate; Feder et al. 2008) to the annual variation of juvenile survival. The timing of parturition is critical to reproductive females, as they must cater to their offspring’s needs in addition to their own. Lactation costs are particularly high in mammals (Sadleir 1982, Clutton-Brock et al. 1989), and could make reproductive females more vulnerable to harsh environmental conditions. Additionally, the timing of parturition can generate reproductive costs for females by lowering the chances of successfully reproducing the next year (Green and Rothstein 1993). In red deer, the later in the year a hind gives birth, the less likely she is to conceive during the following breeding season (Clutton-Brock et al. 1983). These selection pressures acting on newborns and mothers are thought to underlie the commonly observed strong seasonality in timing of birth across ungulates in temperate ecosystems (Bronson 1989).

Although less predictable than temperate and boreal ecosystems, arid and semi-arid ecosystems also show a pronounced seasonality, i.e. wet and dry seasons, mainly driven by the distribution of rains during the year (Owen-Smith and Ogutu 2013). Some large herbivore species give birth within a relatively short time window in such environments (e.g. 80% of birth occur within three weeks in the blue wildebeest (*Connochaetes taurinus*); Sinclair et al. 2000), while others have extended breeding periods (Owen-Smith and Ogutu 2013), or even breed year-round (e.g. in the plains zebra (*Equus quagga*); Sinclair et al. 2000). In such species, and even if a peak of births is often observed during the wet season, some females therefore give birth in less favourable conditions than others (Boutton et al. 1988a, Boutton et al. 1988b, Feng et al. 2013), potentially exposing themselves and their offspring to a higher risk of mortality (Valeix et al. 2009). An interesting and yet unanswered question is: for such species, to what extent does giving birth in the dry “unfavourable” season, reduce the probability of survival of both the newborn and the mother? To better understand the phenology of reproduction and the factors that may influence it, more empirical studies exploring juvenile survival and reproductive costs, both in species reproducing year-round and in tropical large herbivores are needed (Clutton-Brock and Sheldon 2010, Burthe et al. 2011, Festa-Bianchet et al. 2017, but see Lee et al. 2017, MacKay et al. 2018).

In this study, we investigated the effects of season of birth on the annual survival of young and mares in wild plains zebras. We explored the impact of the duration of the dry season, which varies according to the timing of rainfall, on the annual survival of foals, yearlings and adult females. We took advantage of a long-term individual-based monitoring program of plains zebras conducted in Hwange National Park (HNP) in Zimbabwe. Although their environment is seasonal (Chamaille-Jammes et al. 2006), plains zebras breed year-round in this ecosystem (Fig. 1), and are exposed to a high inter-annual variability in the duration of the dry season. This constitutes an adequate framework to study the impact of the timing of reproduction on the survival of young and females in seasonal tropical environments.

**Figure 1:**
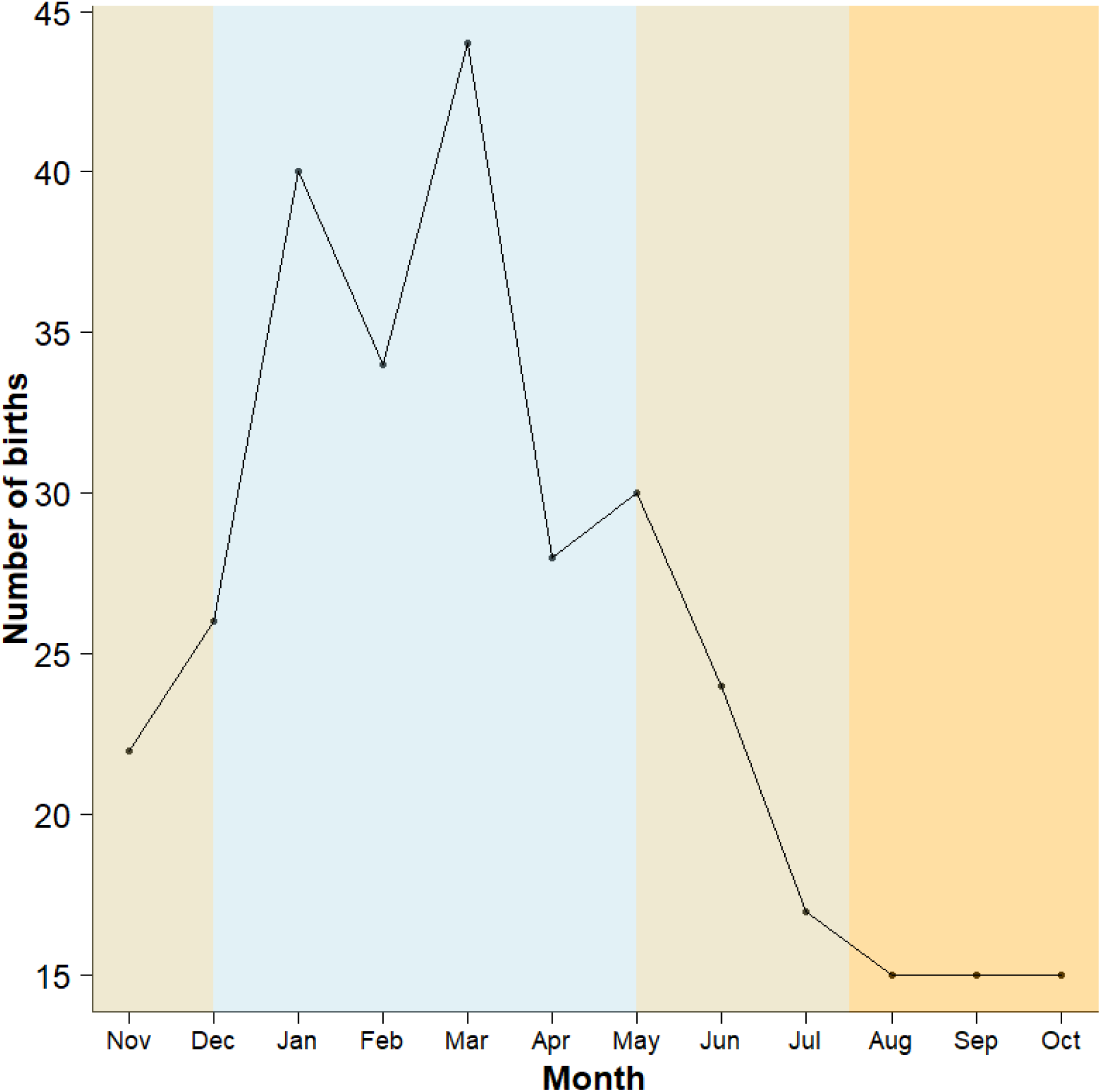
Distribution of plains zebra (*Equus quagga*) births per month, estimated from the individual-based monitoring conducted in Hwange National Park, Zimbabwe, between 2008 and 2019 (*n* = 310 foals). The black line represents the total number of births per month over the complete study period (i.e. all years summed). The coloured areas indicate the seasons: blue = main wet season (approx. mid-December to mid-May), orange = main dry season (approx. August to October), mixed colour = inter-annual variability in the start of the seasons (start of the wet season: November to mid-December; start of the dry season: mid-May to July).

We hypothesised that the dry season, and in particular long dry seasons which lead to harsh conditions for herbivores, should decrease early juvenile survival in plains zebra (i.e. until weaning, between birth and six months of age) because this period corresponds to the most vulnerable stage of life (Caughley 1966, Festa-Bianchet 1988). Similarly, harsh environmental conditions may reduce late juvenile survival, when foals become independent from their mother (between six and 12 months of age). These older foals are arguably less vulnerable but are still growing and cannot rely on their mother anymore for food provisioning and protection (Smuts 1976, Côté and Festa-Bianchet 2001). We expected a comparatively smaller effect of exposure to harsh conditions on yearling survival (one to two years of age) than on foal survival, because older individuals should have more resources to survive through periods of low food availability (Gaillard et al. 2000). Regarding mares, we expected that the costs of reproduction would result in a lower survival rate of reproductive females compared to non-reproductive females. Alternatively, one could expect the opposite, with reproductive females coping better than non-reproductive females. Non-reproductive females could be of lower phenotypic quality, explaining their inability to breed (Reznick et al. 2000, Hamel et al. 2009) and vulnerability to harsh environmental conditions (Weladji et al. 2008).

## Materials and methods

### 1. Context of the study

#### 1.1. Study site

HNP (19°00’ S, 26°30’ E), Zimbabwe, covers 14,651 km^2^ of bushlands and woodlands interspersed with grasslands (Rogers et al. 1993, Arraut et al. 2018). Dominant grass species in HNP are *Aristida sp., Bothriochloa radicans, Cynodon dactylon, Digitaria eriantha, Eragrostis sp., Heteropogon contortus, Panicum maximum, Pogonarthria squarrosa and Schimidtia pappophoroides* (unpub. data). The altitude varies between 900 and 1,100m a.s.l.. Average annual rainfall is approximately 600mm, with marked inter-annual variations (Chamaille-Jammes et al. 2006). On average, 98% of the rainfall events occur during the wet season, generally from October to April (Chamaille-Jammes et al. 2006, Chamaille-Jammes et al. 2007). The start of the rains is characterised by a high inter-annual variability, leading to a variable duration of the wet and dry seasons. The study took place in the north-east of the park, where artificial waterholes retain water year-round and areas that are > 8km from a waterhole are rare, even in the dry season (Chamaille-Jammes et al. 2007). The densities of lion (*Panthera leo*) and spotted hyena (*Crocuta crocuta*), the two main predators of plains zebras, are relatively high (Loveridge et al. 2007, Drouet-Hoguet 2007). There is no hunting in the park.

#### 1.2. Study species

Plains zebras live in harems composed of a stallion, several mares and their offspring under two years of age (Klingel 1969). Females can give birth to a singleton at any time of the year in most of their geographic range, including Zimbabwe (Dasmann and Mossman 1962, Barnier et al. 2012), even if a birth peak occurs from January to March in this area (Fig. 1). Gestation length ranges between 340 and 390 days and foals are weaned between six and 12 months of age (Smuts 1976, Owen-Smith and Mason 2005, Pluháček et al. 2007). Plains zebra foals are considered as “followers” on the hider-follower gradient (Lent 1974), as they stand and follow their mother a couple of hours after birth (Sinclair2000). Plains zebras are grazers (*sensu* Hofman 1989), feeding virtually only on grasses, hence the availability of their food resource is directly linked to rainfall amount and distribution within the year, as well as with the general availability of habitats with a grass layer (Du Plessis 2001). Similarly to the other large herbivores occupying African savannas, the plains zebra is considered to be a capital breeder (Jönsson 1997, Stephens et al. 2009), although there is currently little empirical assessment of its energy allocation strategy (but see Ogutu et al. 2014). In HNP, the population of plains zebras is mostly resident thanks to the artificial waterholes providing water year round inside the park. However some seasonal shifts in home-ranges have been observed for some harems (unpublished GPS data).

#### 1.3. Seasonality and plains zebra life histories

Following the protocol presented in Grange et al. (2015), we recorded the presence of individually identified plains zebras between 2008 and 2019 using visual identification of their unique stripe pattern. Observations were conducted twice a year, around March and around August, during field sessions (hereafter called “sessions”, *n* = 24 sessions, *mean session duration ± SD* = 45 ± 25 days, *range* = 13-88 days). Individuals of more than 24 months were identified as adults while the age of the young individuals was determined more accurately based on their relative size compared to adults, using the criteria of Smuts (1975) and Penzhorn (1982), and photographs of individuals of known age from HNP. We also attributed an accuracy to this age estimation, based on our experience from the field. The age of very young foals was considered accurate to a few days, whereas young seen at an older stage for the first time could be attributed an age ± 2 months, for instance.

The main goal of this study was to explore the effect of seasonality as a categorical (i.e. wet season versus dry season) or continuous (i.e. duration of the dry season) variable on the survival of young and mares, accounting for the reproductive state of the mare and the age-class of the juvenile. We also assessed the probability of transition between reproductive and non-reproductive states in mares from one session to the next. Explanatory variables are described in the following sections and summarised in Table 1.

**Table 1:** Set of explanatory variables used in the models investigating survival in young individuals and mares plains zebras (*Equus quagga*), and transition probability between reproductive and non-reproductive states in mares plains zebras (Hwange National Park, Zimbabwe, 2008-2019).

| Variable | Type | Description | Used in the model(s) |
| --- | --- | --- | --- |
| *Season* | Categorical | “Wet” or “Dry” season | Young & Mares |
| *Time in dry season* | Continuous | Proportion of dry season between two consecutive sessions [range: 9-80 %] | Young & Mares |
| *Time* | Categorical | ID number of the field session [range: 1-24] | Young & Mares |
| *Trap-dependence* | Categorical | The individual has been seen during the previous field session (1) or not (0) | Young & Mares |
| *Age-class* | Categorical | “Younger foal” (less than six months of age), “Older foal” (between six and 12 months of age) or “Yearling” (between one and two years of age) | Young |
| *Reproductive state* | Categorical | The mare is seen with a dependent juvenile (1) or not (0) | Mares |
| *Sight* | Categorical | The individual is seen for the first time (1) or not (0) | Mares |

### 2. Explanatory variables

For each year, we identified the transition date between wet and dry season using 500m spatial resolution, bi-monthly Normalised Difference Vegetation Index (NDVI) records from the NASA website (MOD13A1 product, computed from atmospherically corrected bi-directional surface reflectance masked for water, clouds, heavy aerosols, and cloud shadows; temporal resolution: 15 days; https://modis.gsfc.nasa.gov) and daily rainfall records from the Climate Hazards Center website (Rainfall Estimates from Rain Gauge and Satellite Observations, https://www.chc.ucsb.edu). During the study period and according to our estimations, the wet season in HNP started between the 1^st^ of November and the 19^th^ of December, while the dry season started between the 9^th^ of May and the 29^th^ of July (Fig. 1, Online Resource 1). We attributed a season to each session according to the date of the session (variable *season*, Table 1). We further estimated the proportion of days of dry season between the first day of the session *s* and the first day of the following session *s+1* (variable *pds*, namely “proportion of dry season”, Table 1). We defined three age-classes among the young: “younger foals” of less than six months of age, “older foals” aged between six and 12 months of age, and “yearlings” between one and two years of age (Gaillard et al. 2000, Table 1).

### 3. Statistical analysis

#### 3.1. Capture-Mark-Recapture framework

We used a Capture-Mark-Recapture (CMR, Lebreton et al. 1992) framework to account for the variability of the probability of recapture (i.e. re-observation) of the individuals of this population (Grange et al. 2015). We considered each session as a discrete event, summarised by its starting date. As the time interval between two successive sessions varied, we used the *time.intervals* argument of the *process.data* function to correct survival estimates for the duration of exposure to mortality risk (*RMark* package, Laake 2013). For this, we calculated the proportion of time elapsed between two successive sessions in relation to a year of 365 days: *Δt_session2-session1_ = (Starting date_session2_ - Starting date_session1_) / 365*. We performed all CMR analyses using the program MARK (www.phidot.org/software/mark) and the R statistical software (version 4.3.1, www.r-project.org), thanks to the package *RMark* (Laake 2013). The goodness-of-fit tests (GOF tests: see below) were conducted with the R package *R2ucare* (Gimenez et al. 2017).

#### 3.2. Survival of the young

Similarly to previous demographic analyses performed on these data (Grange et al. 2015, Vitet et al. 2020, Vitet et al. 2021), we built capture histories for each individual as follows: 0 corresponding to “no sighting of the individual during the session”, and 1 corresponding to “at least one sighting of the individual during the session”. We ran the analyses on individuals whose date of birth was known with an accuracy ranging from 0 to ± 90 days, based either on direct observation in the field or on hormonal assay of the faeces of the female (*n* = 310). Indeed, the dataset contained individuals observed at least once in the field (*n* = 290) and individuals never observed in the field but whose mother was detected to be pregnant (*n* = 20). For the latter, the date of parturition was estimated based on opportunistic faecal sampling from the mare and subsequent 20-oxopregnanes and oestrogens enzyme immunoassay (Online Resource 2, Ncube et al. 2011). We recorded these individuals as being identified at birth only and never seen again, assuming no foetal loss or abortion as usually reported in large mammals. Including these individuals in our analysis allowed us to minimise potential survivorship bias that could arise from the fact that field sessions are spaced by more or less six months, a period during which some foals may be born and die after birth without being observed. Not accounting for these unobserved early mortalities could lead to inflated survival probability as only the surviving foals would be recorded.

Individuals with an estimated date of birth falling within a three months period before or after the closest session were assigned to the same cohort and we assumed that they experienced similar environmental conditions during the last months of gestation and beginning of lactation (i.e. *season* and *proportion of dry season* variables). Based on the abundance of literature on northern ungulates breeding synchronously during a restricted period, a cohort is generally defined as the ensemble of individuals born during the same year (Gaillard et al. 1993, Gaillard et al. 2003). However, working on annual cohorts is less meaningful in the case of asynchronous breeders, for which births are more spread out and can happen during different seasons. In our study, it made more sense to define a cohort at the seasonal scale, such as Lee et al. (2017) did to study juvenile survival of giraffes *Girafa camelopardalis*. We kept sub-adults and adults (> two years of age) of known date of birth in the model to obtain better estimations of juvenile and yearling survival. In other words, individuals re-observed at an age of two years or more but missed during the previous sessions still contributed to the estimation of survival of the young, even if adult survival *per se* was the focus of another analysis. Indeed, because detection of individuals is imperfect, a zebra can remain unseen for several years, even if still alive. If a foal is only seen again at the adult stage but adult stage is not included in the juvenile survival analyses, these individuals would be considered as dead and juvenile survival would be overall biased low.

In a preliminary step, we performed goodness-of-fit tests to check if the dataset of young individuals did fit the CMR model assumptions (Lebreton et al. 1992). The GOF tests of the fully time-dependent model (Gimenez et al. 2018) highlighted possible overdispersion (Test 2.CL: *χ^2^* = 40.79, *df* = 16, *P* < 0.01; Test 3.Sm: *χ^2^* = 11.05, *df* = 19, *P* = 0.92), owing to trap-dependence (*χ^2^* = 139.35, *df* = 17, *P* < 0.01) and transience issues (*χ^2^* = 70.47, *df* = 22, *P* < 0.01). After examination of Test 2.CL, we noticed that overdispersion was mainly caused by three sessions over the 24 available in the dataset, therefore we considered this lack of fit as marginal and ignored overdispersion in the analyses.

We accounted for trap-dependence by adding a trap-dependence effect in the recapture model (*td* variable, Table 1). As the distribution of plains zebras within the park could depend on seasonal dynamics, we tested the effect of the *season* or *proportion of dry season* in the recapture model. Transience was likely due to the age structure, as young individuals often have lower survival in large herbivores (Gaillard et al. 2000) and are recaptured at a lower frequency than adults, leading to a lower recapture probability. Thus, we fitted the survival model with an *age-class* effect (Table 1). We also explored the effect of several poolings of those age-classes (e.g. older foals and yearlings pooled together as one single age-class versus younger foals). This was done for methodological reasons: pooling several age classes that do not behave significantly different from one another allows for a more accurate and robust estimation by increasing the sample size used in the models. We tested the effect of *season* or *proportion of dry season* in the young survival models (Online Resource 3). The combination of these different environmental and age class parameters led to the comparison of a set of 22 survival models (Online Resource 3). For comparison purposes, we also conducted an analysis of juvenile survival using a Generalised Linear Model (GLM) instead of a CMR approach (Online Resource 4).

#### 3.3. Mare survival

To study the probability of survival and transition between reproductive states in adult females (*n* = 322), we used a multi-state capture-recapture model (Lebreton et al. 2009). We built capture histories for each individual as follows: 0 corresponding to “no sighting of the mare during the session”, 1 corresponding to “all sightings of the mare without any dependent juvenile during the session” and 2 corresponding to “at least one sighting of the mare with a dependent juvenile during the session”. We considered a dependent juvenile (corresponding to our “younger foal” age-class) as an individual that was still suckling and entirely relying on its mother’s milk to feed, so under six months of age. In CMR analyses, death and emigration might be confounded as they both result in the disappearance of the individual from the system. Although zebras are mostly philopatric in HNP (pers. obs.), it is possible that a few individuals move in and out the study site at times. CMR estimates of survival in our study thus correspond to “apparent survival” confounding permanent emigration and death (see Lebreton et al. 1992, Pollock 1981). For simplicity and following common practice, we nevertheless refer to adult female survival in the following.

As there were too few observations to conduct the GOF tests of the multi-states model, we conducted the GOF tests on the single-state model instead (i.e. without considering reproductive states). The tests revealed overdispersion (Test 2.CL: *χ^2^* = 49.02, *df* = 19, *P* < 0.01; Test 3.Sm: *χ^2^* = 72.10, *df* = 20, *P* < 0.01), trap-dependence (*χ^2^* = 112.13, *df* = 21, *P* < 0.01) and transience (*χ^2^* = 62.23, *df* = 21, *P* < 0.01). After examination of Test 2.CL and 3.Sm, we noticed that overdispersion was mainly caused by five and three sessions over the 24 available in the dataset, respectively. Again, we considered it as marginal and ignored overdispersion in the analyses. We accounted for trap-dependence by adding a trap-dependence effect (*td* variable, Table 1) in the recapture model. To account for transience, we added a categorical variable in the survival model contrasting mares observed for the first time during the survey and mares previously captured at least once during the survey, following the method described by Pradel et al. (1997; *sight* variable, Table 1). We evaluated the effect of *reproductive state* (Table 1), and *season* or *proportion of dry season* on recapture, survival and transition probabilities. Age being not known for most adult females, we were unable to include this variable in the model. We compared a set of 12 survival models and 11 transition models (Online Resource 3).

#### 3.4. Model selection

For both young and mares, we conducted model selection by comparing models ranging from the most complex full time-dependent model (*time* variable, i.e. chronological number identifying the session, Table 1) to the reference model (i.e. depending exclusively on the covariables related to GOF corrections, Online Resource 3). Because of the large number of possible combinations between recapture and survival (and transition for the mares dataset) models, we conducted a sequential model selection. We started by modelling recapture, then survival and finally transition probabilities (Lebreton et al. 1992). We conducted a first model selection to identify the most parsimonious recapture model, while the survival model was set to depend exclusively on the covariables related to GOF corrections (i.e. *age-class* for young individuals and *sight* for mares) and transition probability held constant for mares. We then modelled survival probability using the best recapture model and constant transition probability for mares. We eventually selected the most parsimonious transition probability model with the most parsimonious recapture and survival models for mares.

We used the Akaike Information Criterion adjusted for small sample sizes (AICc) and selected the model with the lowest number of parameters for models having a ΔAICc < 2 (i.e. models receiving substantial empirical support, Burnham and Anderson 2002, Arnold 2010) when compared to the top model (parsimony principle, Burnham and Anderson 2002). We reported the number of models within a ΔAIC < 7 (i.e. models receiving substantial or some form of empirical support, Burnham and Anderson 2002, Arnold 2010) from the top model which included the effect of the *proportion of dry season* or the *season* on the probability of survival. Following Arnold (2010), we reported the 85% confidence intervals of estimated parameters, in accordance with our AICc model selection procedure. We used the scaled value of *proportion of dry season* in the models to ease model convergence, but for convenience, we back-transformed the coefficients to their initial scale in the Results section.

## Results

### 1. Survival of the young individuals

Results on recapture probabilities of juveniles are presented in Online Resource 5. In summary, recapture probability was best described by an effect of *trap dependence* and *proportion of dry season*. Recapture probability ranged between 0.293 ± 0.039 SE, 85% CI [0.240; 0.353] and 0.483 ± 0.030 SE, 85% CI [0.440; 0.526] over the study period. According to the most parsimonious model, survival varied between age-classes (see Online Resource 3 for the detailed model selection). Early juvenile survival was lower (*φ* = 0.458 ± 0.044 SE, 85% CI [0.395; 0.522]) than late juvenile and yearling survival. Additionally, late juvenile and yearling survival decreased with the *proportion of dry season* (*β* = −0.026 ± 0.016 SE, 85% CI [−0.049; −0.002]). The late juvenile and yearling survival (older foals and yearlings pooled together in the most parsimonious model) varied from 0.850 ± 0.095 SE, 85% CI [0.661; 0.943] when the *proportion of dry season* was the smallest (i.e. 9 % of the time) to 0.480 ± 0.120 SE, 85% CI [0.316; 0.648] when the *proportion of dry season* was the largest (i.e. 80 % of the time, Fig. 2). The *season* variable was not retained in the most parsimonious models of either the CMR or the GLM approaches (Online Resource 4). Although not significant, late juvenile and yearling survival was lower and more variable during the dry (0.536 ± 0.221 SD) than during the wet season (0.707 ± 0.172 SD). Overall, the effect of the *proportion of dry season* on the probability of survival was present in *n* = 7 models in a ΔAIC < 7, whereas the effect of the *season* was present in *n* = 6 models with ΔAIC < 7.

**Figure 2:**
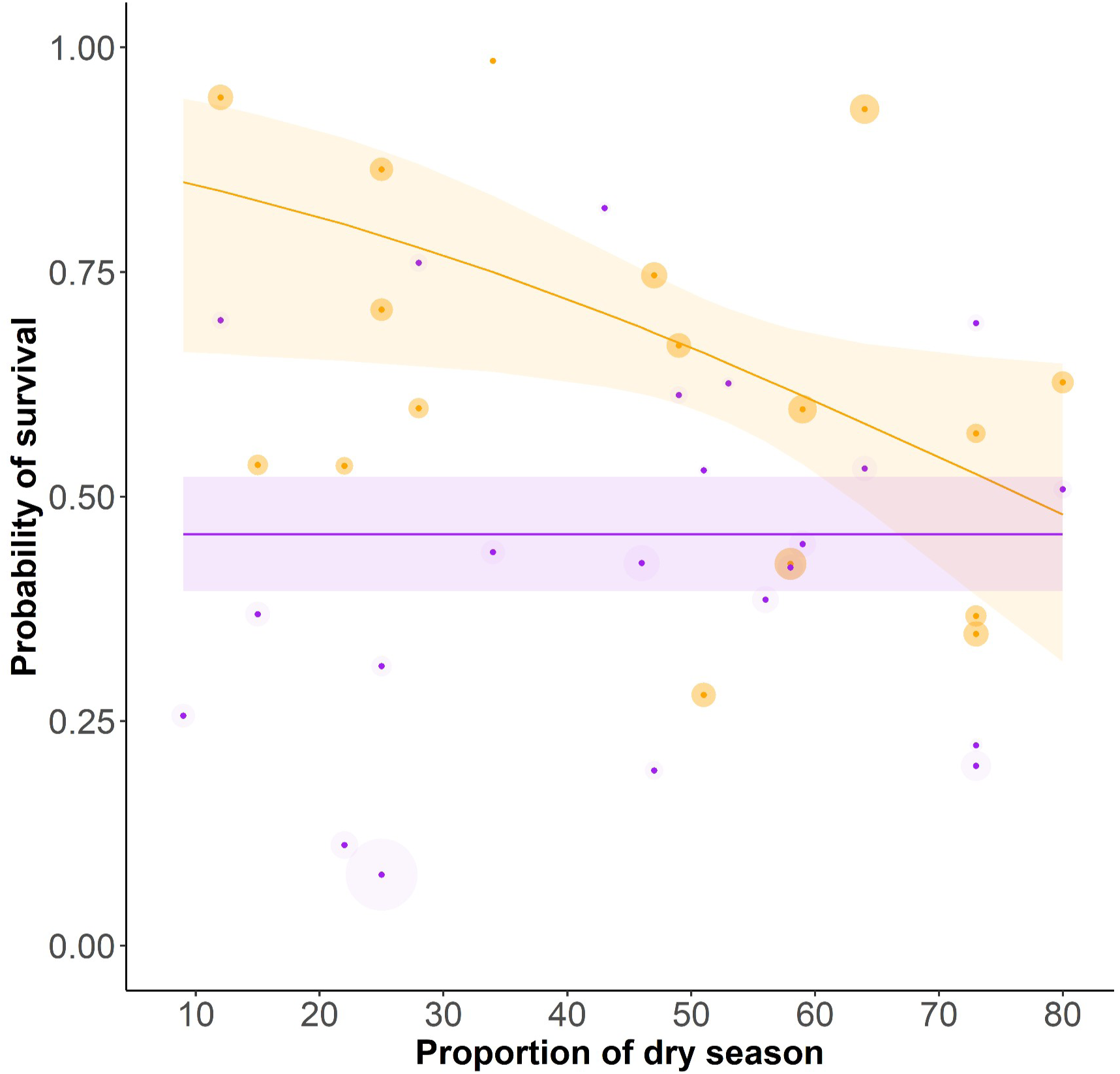
Probability of annual survival of plains zebra (*Equus quagga*) young individuals in Hwange National Park, Zimbabwe (2008-2019), according to the proportion of dry season between two consecutive sessions. Purple: early juvenile survival (younger foals, between birth and six months of age), orange: late juvenile and yearling survival (older foals, between six and 12 months of age and yearlings, between one and two years of age). Solid lines represent predicted values from the top model. Shaded areas represent 85% confidence intervals of these predicted values. Dark coloured dots represent the survival predicted by the time model, shaded areas around the dot are proportional to 1/standard error of these predicted values.

### 2. Mare survival

Recapture probability was higher for mares than for young individuals, within the range 0.449 ± 0.033 SE, 85% CI [0.402; 0.496] - 0.849 ± 0.083 SE, 85% CI [0.690; 0.934] (Online Resource 5). The most parsimonious model for survival probability included an additive effect of *sight*, *proportion of dry season* and *reproductive state*. It further included an additive effect of *time* and *reproductive state* for transition probability (see Online Resource 3 for the detailed model selection). As the *sight* variable had a significant effect on the probability of survival, we presented results for mares in their second and following observations only. Results relying on mares observed multiple times serve as a better representation of the average behaviour of the population than those from mares observed for the first time or only once, as the latter could be transient young adult females. The *proportion of dry season* had a negative effect on survival (*β* = −0.035 ± 0.011 SE, 85% CI [−0.050; −0.019], Fig. 3a). The probability of survival of reproductive females varied slightly from 0.995 ± 0.006 SE, 85% CI [0.970; 0.999] when the *proportion of dry season* was the smallest to 0.945 ± 0.059 SE, 85% CI [0.768; 0.989] when the *proportion of dry season* was the largest. Although the interaction between the *proportion of dry season* and the reproductive state of the females was not retained in the top model, we noted that the probability of survival of non-reproductive females appeared to vary more largely, from 0.966 ± 0.019 SE, 85% CI [0.924; 0.985] to 0.703 ± 0.057 SE, 85% CI [0.615; 0.779] as the *proportion of dry season* increased. Overall, the effect of the *proportion of dry season* on the probability of survival was present in *n* = 3 models in a ΔAIC < 7, whereas the effect of the *season* was present in *n* = 0 models in a ΔAIC < 7.

**Figure 3:**
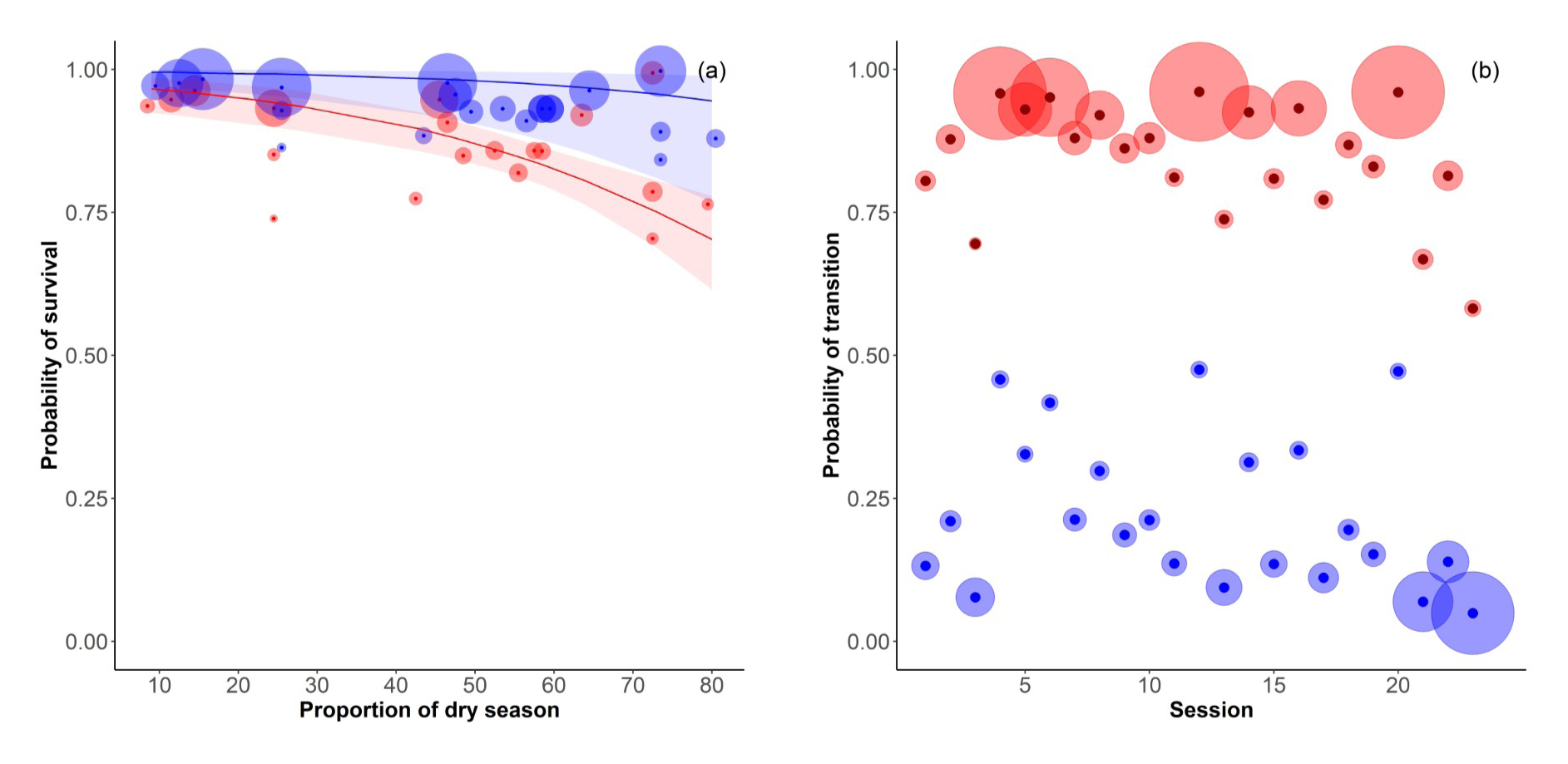
a) Probability of annual survival of plains zebra (*Equus quagga*) mares in Hwange National Park, Zimbabwe (2008-2019), according to the proportion of dry season between two consecutive sessions and the reproductive state (for mares in their second and subsequent observations only, see text for details). Red: non-reproductive females, blue: reproductive females. Solid lines represent predicted values from the top model. Shaded areas represent 85% confidence intervals of these predicted values. Dots represent females survival predicted by the time model, vertical bars represent 85% confidence intervals of these predicted values. b) Probability of transition between reproductive and non-reproductive states for plains zebra mares according to the session. Red: from reproductive to non-reproductive state, blue: from non-reproductive to reproductive state. Dark coloured dots represent the survival predicted by the time model, shaded areas around the dot are proportional to 1/standard error of these predicted values.

The probability of transition between reproductive states was not dependent on either the *season* or the *proportion of dry season* as none of these variables were selected in the top model, but was highly variable according to *time*. The probability for a mare to switch from the reproductive to the non-reproductive state between two consecutive sessions was higher (from 0.582 ± 0.094 SE, 85% CI [0.443; 0.709] to 0.961 ± 0.015 SE, 85% CI [0.933; 0.978]) than the probability to transition from the non-reproductive to the reproductive state (from 0.049 ± 0.018 SE, 85% CI [0.028; 0.083] to 0.475 ± 0.093 SE, 85% CI [0.346; 0.608], Fig. 3b). The probability to be reproductive in the year *n+1* was not different between reproductive females (0.423) and non-reproductive females (0.420, estimates based on the model including the season and reproductive state as explanatory variables).

## Discussion

Our study highlighted an influence of the duration of the dry season on survival of some juvenile age-classes. We found the year-round breeding strategy of female plains zebras to have little consequences on early juvenile survival, which could be related to the capital breeding strategy of energy allocation to reproduction. We did however document variability both within and between years in late juvenile and yearling survival rates with potential consequences for population dynamics.

We did not find major evidence for an effect of the season as a categorical variable, possibly because our field sessions sometimes happened at different stages of a given season due to variability in their timing. However, we identified a marked effect of the duration of the dry season, which better captures the inter-annual variability in environmental seasonality due to its continuous nature. In contrast with early juvenile survival, late juvenile and yearling survival was lower during long dry seasons. Sensitivity to environmental conditions of young individuals is a commonly observed pattern, even in tropical herbivores. For instance, juvenile and yearling survival is highly correlated with the preceding annual rainfall in greater kudus (*Tragelaphus strepsiceros*, Owen-Smith 1990) and calf survival decreases during the dry season in wildebeests (Mduma et al. 1999). Georgiadis et al. (2003) suggest that plains zebra foals under one year of age are more sensitive to fluctuations in rainfall than adults. Young plains zebras only reach their adult size around three years of age and adult body mass around five years of age (Smuts 1975). During this critical period of early growth and development, older foals and yearlings showed a decreasing survival with increasing time spent in dry season. This decrease in survival could result from energy allocation to body growth that may induce a higher vulnerability to poor environmental conditions or predation (Grange et al. 2004).

Reproductive females usually time their reproduction to coincide with the most costly part of the reproductive cycle, corresponding to late gestation and early lactation in large herbivores (Clutton-Brock et al. 1989), and overlap with the period of best food quality or availability (Bronson 1989). In mammals, this matching between food resources and costly reproductive events should minimise subsequent fitness costs such as a lower survival, litter size or probability of reproducing again at the following opportunity. Accordingly, most of the plains zebra mares in HNP gave birth during the wet season. Our results suggest that when environmental conditions are harsher (i.e. longer dry seasons), mothers appeared to buffer the costs for the dependent foal (i.e. younger foal). This supports the idea that plains zebras might be capital breeders, with females engaging in reproduction conditional to a sufficient acquisition of energy prior to the reproductive period. This would increase their ability to cope with their offspring’s requirements in case of an unexpectedly long dry season. However, dry season births do not imply death for either the foal or the mare. Plains zebras can survive on rather low quality grasses, even if their body condition is lower in the dry season (Duncan et al. 2024). Therefore, reproductive timing leading to dry season births may not be strongly selected against. Additionally, the presence of permanent waterholes in the study area reduces the effect that droughts usually have on herbivores via reduced water availability.

Given our results and somewhat counter-intuitively, the optimal timing of births in order to minimise mortality risks of young plains zebras should be the beginning of the dry season. Juveniles could thus benefit from their mother’s protection during the harsher season and enter the age of independence during the most favourable season. Similarly, Lee et al. (2017) showed that giraffes born during the dry season experience a higher survival probability in Tanzania. Although most of the plains zebras’ births in our population happened during the wet season, ensuring sufficient food resources for the mother at a critical stage of their reproductive cycle, the birth peak also extended towards the beginning of the dry season, which might be a more favourable time for the offspring. Such a phenomenon could illustrate a potential trade-off between minimising mother’s reproductive costs and maximising juveniles’ survival, as described in primates (Dezeure et al. 2021).

The length of the dry season only had a small impact on the survival of reproductive females, whose survival always remained very high (∼ 0.95 or above). Our results also suggest that reproductive females were not more sensitive, and possibly were less sensitive, than non-reproductive females to environmental conditions. Females engaging in reproduction thus appeared to not only be able to buffer the effect of the environment on their young foal, but also to bear the costs of reproduction, even under harsh conditions, with minimal impact. As shown by the similar probability of transition to the reproductive state, both from the non-reproductive and the reproductive state the previous year (0.420 and 0.423 respectively), reproductive females were as able as non-reproductive females in engaging in future reproduction. All these results point toward the fact that reproductive females might be those of higher reproductive potential, or experience. If these effects have been well described and studied in temperate or boreal ungulates (e.g. Hamel et al. 2009), our results suggest these observations may apply to some tropical species as well. As both reproductive and non-reproductive females can be found in the same harems, therefore with similar landscape use, we can rule out differences in habitat availability or habitat selection strategies explaining the observed contrasts between females of different reproductive states. Given the importance of age and dominance status, which are often related in equids, on reproduction as previously demonstrated in horses and zebra species (Lloyd and Rasa 1989, Pluháček et al. 2007, Rubenstein and Nuñez 2009), we suspect that they are key traits underlying the observed difference. Unfortunately, age, which was sometimes available (dominance status was not) could not be included in our analysis without reducing its power.

Overall, we documented variations in demographic rates (young survival and female reproductive rates) of plains zebras in HNP, both between seasons and across the years, partly accounted for by the duration of the dry season. Such variations have not been extensively studied in tropical ecosystems so far (e.g. in greater kudus, Owen-Smith 1990). The seasonal variation of juvenile survival has the potential to generate short-term and long-term effects on demographic rates (Gaillard et al. 2003). Compared to strictly seasonal breeders, plains zebras have more variable recruitment rates (Owen-Smith and Mason 2005). Because population dynamics is sensitive to temporal variance in demographic rates, a large variability in juvenile survival might reduce long-term stochastic population growth rate (Tuljapurkar 1989).

## Acknowledgments

This long-term demographic survey was initiated by Patrick Duncan, who also took part in many of the field sessions. The authors are therefore indebted to him, and also acknowledge all the other people involved in data collection over the years, in particular S. Grange, F. Barnier, H. Ncube, C. Vitet and A. Duncan. J. Dezeure assisted with NDVI and rainfall data retrieval. The authors appreciate the insightful comments and proofreading during manuscript preparation by Nicholas van Rooyen. The authors thank Marco Festa-Bianchet and an anonymous reviewer for insightful comments on previous drafts.

## Declarations

### Ethical approval

Ethics approval was not required for this study according to local legislation [Parks and Wildlife Act].

### Consent to participate

Not applicable.

### Consent for publication

Not applicable.

### Availability of data and material

Data available from the corresponding author on reasonable request.

### Code availability

Not applicable.

### Authors’ contributions

LT, CB and SCJ were involved in the Conceptualization, Investigation, Methodology, Writing, Funding acquisition. CB and SCJ were involved in the Supervision and Validation. LT and CB were involved in the Formal analysis. LT and SCJ were involved in the Data curation. LT was involved in the Visualization. SCJ was involved in the Data collection, Project administration.

