## supporting information for "Good moms: dependent-young and their mothers cope better than others with longer dry season in plains zebras"

### Electronic Supplemental Material

##### Author details

[1] Laboratoire de Biométrie et Biologie Évolutive, Unité Mixte de Recherche 5558, Bâtiment 711, Université Lyon I, 43 Boulevard du 11 novembre 1918, F-69622 Villeurbanne Cedex, France.

[2] CEFE, Université Montpellier, CNRS, EPHE, IRD, Montpellier, France.

[3] Mammal Research Institute, Department of Zoology & Entomology, University of Pretoria, Pretoria, South Africa.

[4] LTSER France, Zone Atelier "Hwange", Hwange National Park, Bag 62, Dete, Zimbabwe-CNRS HERD (Hwange Environmental Research Development) program.

**Online Resource 1**: Determination of the transition dates between wet and dry seasons in Hwange National Park, Zimbabwe, during the study period (2008-2019) using NDVI and rainfall records.

We extracted the mean NDVI and rainfall values for the study area using Google Earth Engine facilities (earthengine.google.com/, Gorelick et al. 2017). We used 500 m resolution bi-monthly Normalised Difference Vegetation Index (NDVI) raster from the NASA website (MOD13A1 product, https://modis.gsfc.nasa.gov) and daily rainfall raster from the Climate Hazards Center website (Rainfall Estimates from Rain Gauge and Satellite Observations, https://www.chc.ucsb.edu) between 2007 and 2020.

Based on the work of Chamaillé-Jammes et al. (2006) and field observations, we considered that January, February and March generally fall during the wet season, and that August, September and October generally fall during the dry season. We used those periods as reference periods to calculate the mean NDVI during wet and dry season ($mean_{NDVI\ Wet}$ = 0.68 ± 0.05 SD, $mean_{NDVI\ \mathrm{Dry}}$ = 0.34 ± 0.03 SD). We calculated the mean of both values ($mean_{NDVI\ Wet\ \&\ Dry}$ = 0.51) to obtain the threshold between dry and wet season. We also evaluated the maximum NDVI that occurs during dry seasons during our study period ($max_{NDVI\ \mathrm{Dry}}$ = 0.42) to use it as the threshold between dry and wet season.

We used the fact that NDVI peaks around August and drops around February every year during our study period to narrow our search window as follows: i) we searched for the first NDVI record after the month of August above $max_{NDVI\ \mathrm{Dry}}$ to determine the transition date between wet and dry season every year; ii) we search for the first NDVI record after the month of February bellow $mean_{NDVI\ Wet\ -\ Dry}$ to determine the transition date between dry and wet season every year. Finally, we visually checked the consistency of our estimations based on NDVI with rainfall records (Fig. S1.1). We noticed that rainfalls occur slightly earlier than our estimations of the beginning of the wet season every year, consistent with the expected latency period before vegetation growth in response to the increase of water availability.

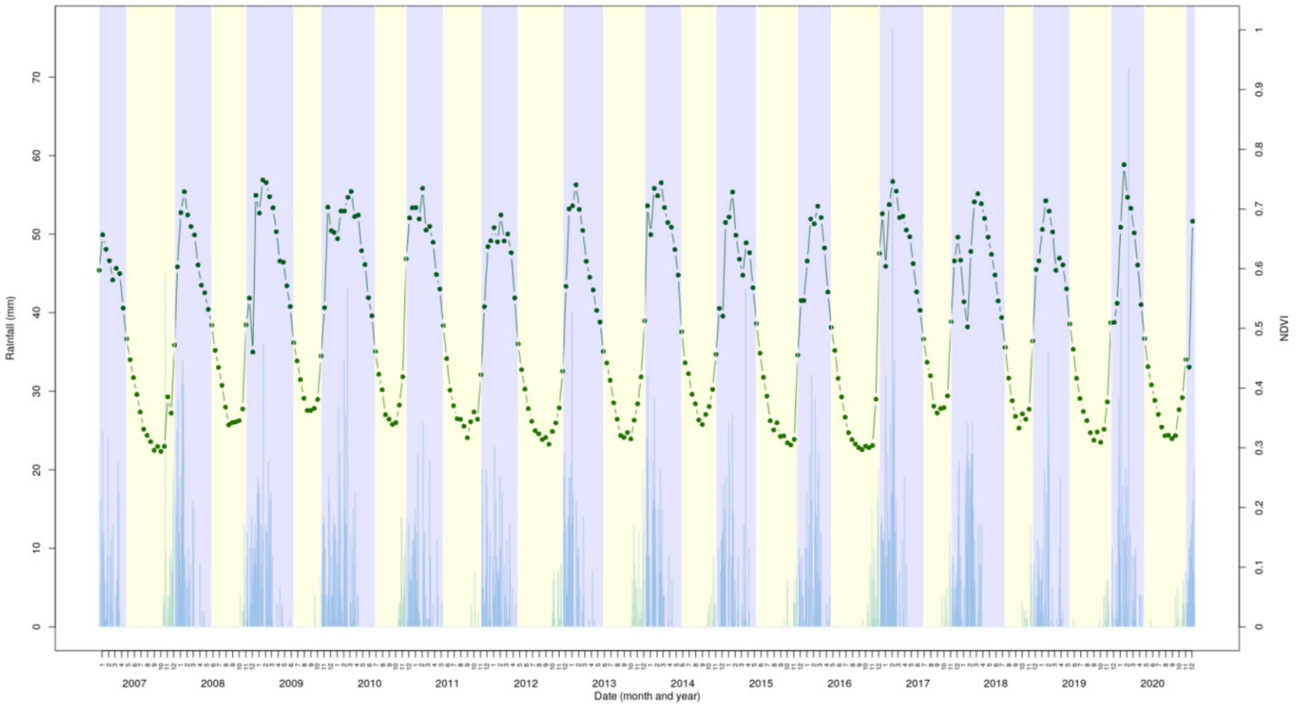


Figure S1.1: mean NDVI and rainfall records in the area of Main Camp, in Hwange National Park, Zimbabwe, between 2007 and 2020. We used these records to estimate the transition dates between wet and dry season (see text for details). Green dots: NDVI bi-monthly records; blue bars: daily rainfall (in mm); blue shade: estimated period of the wet season; yellow shade: estimated period of the dry season.

**Online Resource 2**: Identification of the date of birth of unseen juveniles using hormonal samples from the mares.

Opportunistic faecal samples (*n* = 556) from female plains zebras collected during sessions between 2007 and 2017 were used in this study. We assessed the gestation stage of each mare based on each hormones sample available according to the level of 20-oxopregnanes (fpm) and oestrogens (fem) and using the following decision table (Table S2.1, see also Ncube et al. 2011):

Table S2.1: decision table to assess gestation according to the level of 20-oxopregnanes (fpm) and oestrogens (fem) in the faeces.

| **Gestation stage** | **Detailed gestation stage** | **Level of fpm (ng/g)** | **Level of fem (ng/g)** |
|---|---|---|---|
| No or early pregnancy | Not informative about parturition date | <30 | <100 |
| Mid pregnancy | 9 to 3 months prior to parturition | <30 | >100 |
| | | >30 | >140 |
| Late pregnancy | ≤ 3 months prior to parturition | >30 | <140 |

We estimated all potential periods (as defined in the “Detailed gestation stage” column of Table S2.1) of parturition according to the samples identifying “late” or “mid” pregnancy only (*n* = 264). We estimated the periods when parturition seemed really unlikely using the samples identifying “no or early” pregnancy for females for which at least one potential period of parturition could be estimated (*n* = 192). We considered that the gestation period lasts 375 days (Ncube et al. 2011), and we added a 15 days uncertainty around our period estimations.

Then, on the one hand, we searched for overlaps between the potential periods of parturition predicted; on the other hand, we searched for overlaps between the periods during which parturition seemed very unlikely. We only kept periods supported by at least two different samples (*n* = 65 and 50 for potential periods of parturition and periods during which parturition seemed very unlikely, respectively).

We finally removed the very unlikely periods of parturition from overlapping potential periods of parturition. We took the middle of the estimated period as date of birth (*DOB*), and the range of the potential period of birth divided by 2 as uncertainty (*Acc*) around the date of birth (*n* = 64).

We checked for redundancy between periods of parturition estimated via the hormone samples and juveniles observed in the field by looking for overlaps between the potential periods of births of those two categories of individuals. For juveniles observed in the field, we defined the potential period of birth as the period spanning the time interval [*DOB-Acc; DOB*+*Acc*]. We found *n* = 28 individuals estimated thanks to hormone samples that were effectively observed in the field and removed them from our dataset of juveniles identified thanks to hormone samples. We finally checked for estimated periods of birth (*via* hormones samples) not overlapping any known potential period of birth (*via* field observations), but happening too close to another date of birth, i.e. in a time interval <375 days (i.e. one gestation length) (*n* = 9). We removed them too from our dataset of juveniles identified thanks to hormone samples. We finally added *n* = 27 unseen juveniles estimated via hormone samples to our dataset, over which *n* = 20 were estimated with an accuracy of maximum ± 90 days on their *DOB* and thus retained in the analyses.

**Young individuals**

1. Recapture p (with survival fixed as Phi~age_class):

| **model** | **number of parameters** | **AICc** | **Deviance** | **Delta AICc** | **weight** |
|---|---|---|---|---|---|
| Phi(~age_class)p(~td + time) | 25 | 1731.602 | 1679.907 | 0 | 0.317 |
| Phi(~age_class)p(~td + age_class + time) | 28 | 1731.887 | 1673.761 | 0.285 | 0.275 |
| Phi(~age_class)p(~td + pds) | 7 | 1732.806 | 1718.663 | 1.204 | 0.173 |
| Phi(~age_class)p(~td + age_class + pds) | 10 | 1733.479 | 1713.198 | 1.877 | 0.124 |
| Phi(~age_class)p(~td + season) | 7 | 1735.142 | 1720.999 | 3.54 | 0.054 |
| Phi(~age_class)p(~td + age_class + season) | 10 | 1736.166 | 1715.885 | 4.564 | 0.032 |
| Phi(~age_class)p(~td + age_class * pds) | 13 | 1738.033 | 1711.566 | 6.431 | 0.013 |
| Phi(~age_class)p(~td) | 6 | 1739.86 | 1727.754 | 8.258 | 0.005 |
| Phi(~age_class)p(~td + age_class) | 9 | 1740.152 | 1721.922 | 8.55 | 0.004 |
| Phi(~age_class)p(~td + age_class * season) | 13 | 1740.792 | 1714.325 | 9.191 | 0.003 |
| Phi(~age_class)p(~td + age_class * time) | 85 | 1797.712 | 1607.033 | 66.11 | 0.000 |
| Phi(~age_class)p(~1) | 5 | 1835.724 | 1226.94 | 104.122 | 0.000 |

Age_class = foal_young+foal_old+yearling+adult; td = trap-dependence, pds = proportion of dry season.

2. Survival Phi (with recapture fixed as p~td + pds):

| model | number of parameters | AICc | Deviance | Delta AICc | weight |
|---|---|---|---|---|---|
| Phi(~pds:foal_old_yearling + foal_young + adult)p(~td + pds) | 7 | 1729.988 | 1715.845 | 0 | 0.301 |
| Phi(~age_class + pds)p(~td + pds) | 8 | 1731.713 | 1715.53 | 1.725 | 0.127 |
| Phi(~pds:foal_old + foal_young + yearling + adult)p(~td + pds) | 8 | 1732.444 | 1716.261 | 2.456 | 0.088 |
| Phi(~-1 + season:foal_old_yearling + foal_young + adult)p(~td + pds) | 7 | 1732.52 | 1718.377 | 2.532 | 0.085 |
| Phi(~age_class)p(~td + pds) | 7 | 1732.806 | 1718.663 | 2.818 | 0.073 |
| Phi(~age_class + season)p(~td + pds) | 8 | 1733.413 | 1717.229 | 3.425 | 0.054 |
| Phi(~-1 + season:foal_old + foal_young + yearling + adult)p(~td + pds) | 8 | 1733.475 | 1717.291 | 3.487 | 0.053 |
| Phi(~pds:yearling + foal_young + foal_old + adult)p(~td + pds) | 8 | 1733.95 | 1717.766 | 3.961 | 0.041 |
| Phi(~-1 + season:foal + yearling + adult)p(~td + pds) | 7 | 1734.248 | 1720.105 | 4.26 | 0.036 |
| Phi(~-1 + season:foal_young + foal_old + yearling + adult)p(~td + pds) | 8 | 1734.359 | 1718.175 | 4.371 | 0.034 |
| Phi(~pds:foal_young + foal_old + yearling + adult)p(~td + pds) | 8 | 1734.718 | 1718.534 | 4.729 | 0.028 |
| Phi(~-1 + season:yearling + foal_young + foal_old + adult)p(~td + pds) | 8 | 1734.847 | 1718.663 | 4.859 | 0.026 |
| Phi(~age_class * pds)p(~td + pds) | 11 | 1735.143 | 1712.805 | 5.155 | 0.023 |
| Phi(~pds:foal + yearling + adult)p(~td + pds) | 7 | 1735.273 | 1721.13 | 5.285 | 0.021 |
| Phi(~age_class * season)p(~td + pds) | 11 | 1738.776 | 1716.438 | 8.788 | 0.004 |
| Phi(~pds:foal_young_foal_old_yearling + adult)p(~td + pds) | 6 | 1739.337 | 1727.23 | 9.349 | 0.003 |

| | | | | | |
|---|---|---|---|---|---|
| Phi(~-1 + season:foal_young_foal_old_yearling + adult)p(~td + pds) | 6 | 1740.049 | 1727.942 | 10.061 | 0.002 |
| Phi(~age_class + time)p(~td + pds) | 28 | 1742.605 | 1684.479 | 12.617 | 0.001 |
| Phi(~-1 + time:foal_young + foal_old + yearling + adult)p(~td + pds) | 28 | 1747.125 | 1688.999 | 17.137 | 0.000 |
| Phi(~-1 + time:foal_young_foal_old_yearling + adult)p(~td + pds) | 24 | 1749.297 | 1699.735 | 19.309 | 0.000 |
| Phi(~-1 + time:foal + yearling + adult)p(~td + pds) | 27 | 1750.441 | 1694.465 | 20.453 | 0.000 |
| Phi(~1)p(~td + pds) | 4 | 1813.841 | 1805.79 | 83.853 | 0.000 |

td = trap-dependence; pds = proportion of dry season; foal = foal_young and foal_old gathered; foal_old_yearling =foal_old and yearling gathered; foal_young_foal_old_yearling = foal_young, foal_old and yearling gathered.

**Mares**

1. Recapture p (with survival fixed as S~sight and transition fixed as Psi~1):

| model | number of parameters | AICc | Deviance | Delta AICc | weight |
|---|---|---|---|---|---|
| S(~sight)p(~td + repro_status_t1 * season)Psi(~1) | 8 | 3365.045 | 3348.914 | 0 | 0.621 |
| S(~sight)p(~td + repro_status_t1 + season)Psi(~1) | 7 | 3366.032 | 3351.93 | 0.987 | 0.379 |
| S(~sight)p(~td + repro_status_t1 + pds)Psi(~1) | 7 | 3388.819 | 3374.717 | 23.774 | 0.000 |
| S(~sight)p(~td + repro_status_t1 * pds)Psi(~1) | 8 | 3389.763 | 3373.632 | 24.717 | 0.000 |
| S(~sight)p(~td + repro_status_t1)Psi(~1) | 6 | 3411.854 | 3399.778 | 46.809 | 0.000 |
| S(~sight)p(~td + repro_status_t1 + time)Psi(~1) | 27 | 3478.092 | 3422.692 | 113.047 | 0.000 |
| S(~sight)p(~td + repro_status_t1 * time)Psi(~1) | 48 | 3499.299 | 3398.858 | 134.254 | 0.000 |
| S(~sight)p(~td + season)Psi(~1) | 6 | 3587.921 | 3575.845 | 222.876 | 0.000 |
| S(~sight)p(~td + pds)Psi(~1) | 6 | 3598.026 | 3585.949 | 232.98 | 0.000 |
| S(~sight)p(~td)Psi(~1) | 5 | 3605.565 | 3595.51 | 240.52 | 0.000 |
| S(~sight)p(~1)Psi(~1) | 4 | 3621.583 | 2566.376 | 256.538 | 0.000 |
| S(~sight)p(~td + time)Psi(~1) | 26 | 3640.988 | 3587.689 | 275.942 | 0.000 |

td = trap-dependence; pds = proportion of dry season; sight = first capture or not for a given female; repro_status_t1 = reproductive state in the current session; repro_status_t2 = reproductive state in the following session.

2. Survival S (with recapture fixed as p~td + repro_status_t1 + season and transition fixed as Psi~1):

| model | number of parameters | AICc | Deviance | Delta AICc | weight |
|---|---|---|---|---|---|
| S(~sight + repro_status_t1 + pds)p(~td + repro_status_t1 + season)Psi(~1) | 9 | 3350.251 | 3332.087 | 0 | 0.607 |
| S(~sight + repro_status_t1 * pds)p(~td + repro_status_t1 + season)Psi(~1) | 10 | 3351.577 | 3331.376 | 1.326 | 0.313 |
| S(~sight + pds)p(~td + repro_status_t1 + season)Psi(~1) | 8 | 3355.496 | 3339.365 | 5.246 | 0.044 |
| S(~sight + repro_status_t1 + season)p(~td + repro_status_t1 + season)Psi(~1) | 9 | 3357.494 | 3339.33 | 7.244 | 0.016 |
| S(~sight + repro_status_t1)p(~td + repro_status_t1 + season)Psi(~1) | 8 | 3358.192 | 3342.061 | 7.942 | 0.011 |
| S(~sight + repro_status_t1 * season)p(~td + repro_status_t1 + season)Psi(~1) | 10 | 3359.358 | 3339.157 | 9.107 | 0.006 |
| S(~sight + season)p(~td + repro_status_t1 + season)Psi(~1) | 8 | 3362.108 | 3345.977 | 11.857 | 0.002 |
| S(~sight + repro_status_t1 + time)p(~td + repro_status_t1 + season)Psi(~1) | 29 | 3363.612 | 3303.998 | 13.361 | 0.001 |
| S(~sight)p(~td + repro_status_t1 + season)Psi(~1) | 7 | 3366.032 | 3351.93 | 15.781 | 0.000 |
| S(~sight + time)p(~td + repro_status_t1 + season)Psi(~1) | 28 | 3370.66 | 3313.155 | 20.409 | 0.000 |
| S(~1)p(~td + repro_status_t1 + season)Psi(~1) | 6 | 3379.785 | 3367.709 | 29.535 | 0.000 |
| S(~sight + repro_status_t1 * time)p(~td + repro_status_t1 + season)Psi(~1) | 50 | 3380.811 | 3275.986 | 30.56 | 0.000 |

td = trap-dependence; pds = proportion of dry season; sight = first capture or not for a given female; repro_status_t1 = reproductive state in the current session; repro_status_t2 = reproductive state in the following session.

3. Transition Psi (with recapture fixed as p~td + repro_status_t1 + season and survival fixed as S~sight + repro_status_t1 + pds):

| model | nb param. | AICc | Deviance | Delta AICc | weight |
|---|---|---|---|---|---|
| S(~sight + repro_status_t1 + pds)p(~td + repro_status_t1 + season)Psi(~time + repro_status_t1:repro_status_t2) | 32 | 3327.145 | 3261.18 | 0 | 0.984 |
| S(~sight + repro_status_t1 + pds)p(~td + repro_status_t1 + season)Psi(~season:repro_status_t1:repro_status_t2) | 12 | 3336.112 | 3311.827 | 8.967 | 0.011 |
| S(~sight + repro_status_t1 + pds)p(~td + repro_status_t1 + season)Psi(~season + repro_status_t1:repro_status_t2) | 11 | 3337.854 | 3315.614 | 10.71 | 0.005 |
| S(~sight + repro_status_t1 + pds)p(~td + repro_status_t1 + season)Psi(~pds + repro_status_t1:repro_status_t2) | 11 | 3348.069 | 3325.828 | 20.924 | 0.000 |
| S(~sight + repro_status_t1 + pds)p(~td + repro_status_t1 + season)Psi(~-1 + repro_status_t1:repro_status_t2) | 10 | 3348.594 | 3328.394 | 21.449 | 0.000 |
| S(~sight + repro_status_t1 + pds)p(~td + repro_status_t1 + season)Psi(~1) | 9 | 3350.251 | 3332.087 | 23.106 | 0.000 |
| S(~sight + repro_status_t1 + pds)p(~td + repro_status_t1 + season)Psi(~pds) | 10 | 3351.014 | 3330.813 | 23.869 | 0.000 |
| S(~sight + repro_status_t1 + pds)p(~td + repro_status_t1 + season)Psi(~season) | 10 | 3351.155 | 3330.955 | 24.01 | 0.000 |
| S(~sight + repro_status_t1 + pds)p(~td + repro_status_t1 + season)Psi(~time:repro_status_t1:repro_status_t2) | 54 | 3352.146 | 3238.505 | 25.001 | 0.000 |
| S(~sight + repro_status_t1 + pds)p(~td + repro_status_t1 + season)Psi(~pds:repro_status_t1:repro_status_t2) | 11 | 3352.645 | 3330.404 | 25.5 | 0.000 |
| S(~sight + repro_status_t1 + pds)p(~td + repro_status_t1 + season)Psi(~time) | 31 | 3359.424 | 3295.581 | 32.28 | 0.000 |

td = trap-dependence; pds = proportion of dry season; sight = first capture or not for a given female; repro_status_t1 = reproductive state in the current session; repro_status_t2 = reproductive state in the following session.

**Online Resource 4**: Generalised Linear Model approach to estimating the survival of plains zebra (*Equus quagga*) juveniles (Hwange National Park, Zimbabwe, 2008-2019).

In addition to the CMR approach, we estimated the survival of juveniles using a different approach based on the assumption that a foal under six months of age (i.e. younger foal in our analyses) is still fully dependent on its mother and cannot survive without its mother. Based on this assumption, we considered each juvenile seen at least once after its date of birth and for which the mother was re-observed at least once between the first observation of her offspring and six months after her date of parturition ($n$ = 37). Even if the results could be less reliable because after six months old, juveniles can sometimes survive without their mother (Smuts 1976), we also included this age-class to estimate the probability of survival for older foals (i.e. between six and 12 months old, $n$ = 126). For both age-classes (modelled as a categorical variable with two categories, i.e. younger foal of less than six months-old and older foal between six and 12 months old), if the mother was seen alone, the juvenile was considered to be dead (0), whereas if the juvenile was also seen during the same field session, it was considered to be alive (1). For juveniles whose mother was seen more than once between the first observation of the juvenile after its date of birth and 12 months after its date of birth, we kept the last observation only, to prevent intra-individual repetitions.

We fitted two logistic regressions to the data, with (assumed) death or survival as response variable, and age-class of the juvenile at the end of the interval considered and the proportion of dry season experienced since birth at the re-observation of the mother (*pds_2*) as predictors. In the first model, we added the interaction between the age-class and *pds_2*, whereas in the second one we only looked at the effect of *pds_2* on older foals. When the juvenile was assumed dead, there was no way to precisely know the duration spent in the dry season before death because this date was unknown. We also included the duration between the first observation of the juvenile and the re-observation of its mother as an additive predictor in both models to account for the fact that death is more likely as time passes.

Both models were less than 2 AIC units apart from each other ($AIC_{age\_class*pds\_2}$ = 195.673 and $AIC_{foal\_young+foal\_old*pds\_2}$ = 194.144). We focused on the model where *pds_2* acts solely on older foals to compare with the CMR approach. We found a significant effect of duration between first observation of the juvenile and the re-observation of its mother ($\beta$ = -0.015 ± 0.003, $p$ < 0.001), but no significant effect of *pds_2* on the survival of older foals ($\beta$ = -0.017 ± 0.014, $p$ = 0.223). Nevertheless, even if the trend was not statistically significant, the probability of survival of older foals tended to decrease with an increased proportion of dry season, similarly to what we found with the CMR approach (Fig. S4.1). Our results were also very similar to what we found with the CMR approach concerning younger foals, as they had a mean survival of 0.374 ± 0.090 SE, 85 % CI [0.255; 0.510] (see Results section).

The lower survival estimated using the GLM approach compared to the estimate from the CMR approach for younger foals (0.374 ± 0.090 SE, 85 % CI [0.255; 0.510] versus 0.458 ± 0.044 SE, 85% CI [0.395; 0.522]) as well as older foals (0.766 ± 0.093 SE, 85% CI [0.608; 0.873] to 0.500 ± 0.136 SE, 85% CI [0.314; 0.686] versus 0.850 ± 0.095 SE, 85% CI [0.661; 0.943] to 0.480 ± 0.120 SE, 85% CI [0.316; 0.648]) could come from the fact that mothers are sometimes seen without their offspring, even if the juvenile is still alive (juvenile seen again later while its mother was seen alone before it reaches 12 months old, from e.g. observations in dense habitats where juveniles can be difficult to spot). However, it seems unlikely as we found only two juveniles in this case in our sample. Besides, the GLM was conducted on a smaller sample as the mother had to be seen at least once between parturition and one year after for the juvenile to be included in this analysis. The effect of the proportion of dry season on older foals survival was less strong with the GLM approach than with the CMR approach, but the overall older foals survival was quite similar (see Results section). Eventually, it is worth noting that yearlings could not be considered in the GLM approach because their survival cannot be assessed according to the re-observations of their mother anymore. This could alter the comparability of the results of the CMR and the GLM approaches, as we pooled older foals and yearlings together to test for the effect of the *proportion of dry season* on survival in the CMR approach. The two age-classes hence contributed to the estimation of the coefficient of the slope linking survival and *proportion of dry season*, which is not the case in the GLM approach.

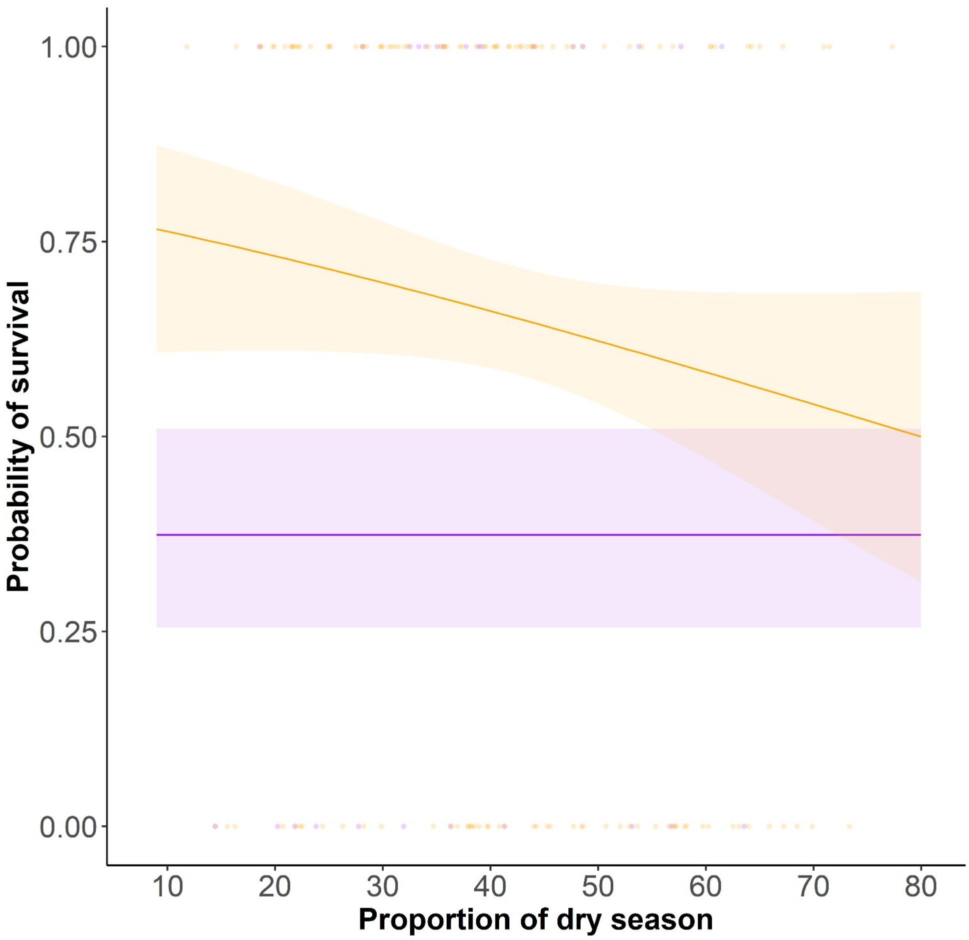


Figure S4-1: Survival according to the proportion of dry season between birth and last re-observation of the mother (before the first year of the juvenile): early juvenile survival (< 6 m.o., purple items, $n$ = 67) and late juvenile survival (> 6 m. o., orange items, $n$ = 126), in plains zebra born in Hwange National Park, Zimbabwe, between 2008 and 2019. For representation purposes, the proportion of dry season was truncated to match the one of the CMR approach, i.e. between 9 and 80 %. The time elapsed between the first observation of the juvenile and the re-observation of its mother was fixed to six months to match the CMR framework. The shaded dots represent the state of the juvenile at the time of the re-observation of its mother (0 = dead, 1 = alive). The solid lines represent the predicted values from the model, for younger (purple) and older (orange) foals. The shaded areas represent the 85% confidence interval (for comparison purposes with the CMR approach) of these predicted values.

**Online Resource 5**: Probabilities of recapture of plains zebra (*Equus quagga*) in young and mares (Hwange National Park, Zimbabwe, 2008-2019).

**Young individuals**

The top model to parsimoniously describe juvenile survival included additive effects of *trap-dependence* and *proportion of dry season* on the recapture probability. The probability of recapture varied from 0.293 ± 0.039 SE, 85% CI [0.240; 0.353] in session 16 (dry season in 2015) to 0.483 ± 0.030 SE, 85% CI [0.440; 0.526] in session 24 (dry season in 2019). The effect of *trap-dependence* on the probability of recapture was positive ($\beta$ = 1.602 ± 0.162 SE, 85% CI [1.368; 1.835]), while the effect of *proportion of dry season* was slightly negative ($\beta$ = -0.013 ± 0.004 SE, 85% CI [-0.018; 0.007]).

**Mares**

The top model included an additive effect of *trap-dependence*, *season* and *reproductive state* on recapture probability. Mean recapture was higher in the wet than in the dry season, and was higher for non-reproductive than for reproductive mares. In the wet season, it varied from 0.829 ± 0.092 SE, 85% CI [0.656; 0.926] to 0.849 ± 0.083 SE, 85% CI [0.690; 0.934] for reproductive mares, and from 0.555 ± 0.037 SE, 85% CI [0.501; 0.608] to 0.590 ± 0.034 SE, 85% CI [0.541; 0.638] for non-reproductive mares. In the dry season, it varied from 0.761 ± 0.127 SE, 85% CI [0.538; 0.897] to 0.792 ± 0.114 SE, 85% CI [0.585; 0.911] for reproductive mares, and from 0.449 ± 0.033 SE, 85% CI [0.402; 0.496] to 0.494 ± 0.027 SE, 85% CI [0.454; 0.533] for non-reproductive mares. There was also a positive effect of *trap-dependence* on the probability of recapture ($\beta$ = 0.661 ± 0.139 SE, 85% CI [0.461; 0.862]).
